# Interpretable Spatial Gradient Analysis for Spatial Transcriptomics Data

**DOI:** 10.1101/2024.03.19.585725

**Authors:** Qingnan Liang, Luisa Solis Soto, Cara Haymaker, Ken Chen

## Abstract

Cellular anatomy and signaling vary across niches, which can induce gradated gene expressions in subpopulations of cells. Such spatial transcriptomic gradient (STG) makes a significant source of intra-tumor heterogeneity and can influence tumor invasion, progression, and response to treatment. Here we report *Local Spatial Gradient Inference* (LSGI), a computational framework that systematically identifies spatial locations with prominent, interpretable STGs from spatial transcriptomic (ST) data. To achieve so, LSGI scrutinizes each sliding window employing non-negative matrix factorization (NMF) combined with linear regression. With LSGI, we demonstrated the identification of spatially proximal yet opposite directed pathway gradients in a glioblastoma dataset. We further applied LSGI to 87 tumor ST datasets reported from nine published studies and identified both pan-cancer and tumor-type specific pathways with gradated expression patterns, such as epithelial mesenchymal transition, MHC complex, and hypoxia. The local gradients were further categorized according to their association to tumor-TME (tumor microenvironment) interface, highlighting the pathways related to spatial transcriptional intratumoral heterogeneity. We conclude that LSGI enables highly interpretable STG analysis which can reveal novel insights in tumor biology from the increasingly reported tumor ST datasets.

## Introduction

Tumor tissues contain heterogeneous cell populations with distinct transcriptional, genetic, and epigenetic features in complex cellular microenvironment^1–3^. Dissecting such multifactorial intratumoral heterogeneity (ITH) is fundamental to understand tumor initiation, metastasis, and therapeutic resistance^4–10^. One source of transcriptional variation in cells is their microenvironments, which shape the gene expression through different ways, such as cell-cell communication (e.g., ligand-receptor signaling) or local signaling cues (e.g., pH, oxygen, metabolites). As a result, some cells would display gradated transcriptional variation along with their spatial localizations, therein termed ‘spatial transcriptomic gradient’ (STG). Identification of STGs can greatly enhance our understanding of spatial-phenotypic relationship of cells, enhancing discovery of multicellular signaling^11^ that are elusive in current cell-type-centric investigations. For instance, oxygen gradient has been shown to shape intra-tumoral heterogeneity affecting tumor proliferation in over 19 tumor types^12–14^.

The development of spatial transcriptomics (ST) technologies^15–17^ allows simultaneous characterization of gene expressions and tissue context of cells in a high-throughput manner, and thus provide sufficient information for systematic identification of STGs in tissue samples. For instance, hallmark pathway gradients have been observed across tumor-TME boundary in liver cancer slides along the pre-defined axis perpendicular to that boundary^18^. However, there is an unmet analytical need to perform *de novo* discovery of STGs without prior pathological annotations, and to discover molecular-spatial heterogeneity beyond apparent pathological annotations. To our best knowledge, no method exists that can detect simultaneously the existence and direction of STGs, which can vary abruptly and substantially across neighboring niches. Trajectory inference (TI) approaches^19,20^ developed for single-cell transcriptomic data analysis cannot be readily applied due to their assumptions on continuity.

Here, we report a novel computational framework, LSGI (Local Spatial Gradient Inference), that performs *de novo* detection, characterization, and visualization of STGs from ST data. LSGI aims at reconstructing salient STGs across spatial niches. As a highly flexible framework, LSGI combines cell phenotype quantification (e.g., pathway activity) with linear models to simultaneously detect the existence and direction of linear spatial gradient in each small niches. It applies NMF to derive quantitative, interpretable cell phenotypes from the gene expression matrix of a ST data. We demonstrated the utility of LSGI in detecting STGs of different cell phenotypes in tumor samples with aberrant cellular composition and tissue reorganization. In particular, we revealed spatial proximity of different phenotypes, highlighting an opposite-directed gradient of neural progenitor-like phenotype and hypoxia phenotype in the intratumoral region of a glioblastoma sample. We further applied LSGI to perform a meta-analysis on 87 publicly available tumor ST datasets from 9 studies. We identified a total of 356 NMF programs associated with STGs and grouped nearly 3/4 of them to 19 meta-programs (MPs). Some of the MPs were shared by multiple tumor types, while others were tumor-type-specific. About 1/4 of the NMF programs were characterized as sample-specific programs, highlighting inter-patient heterogeneity. We further categorized the programs based on their spatial association with tumor region, normal region or boundary regions and highlighted NFKB-TNFA signaling pathways as recurrent gradated programs associated with spatial ITH in different glioblastoma samples, which has been reported as a mechanism employed by GBM cells to enhance their resistance^21^. All the processed data of this meta-analysis have been made publicly accessible (https://zenodo.org/records/10626940), and we provide R code to assist visualization and interpretation of the phenotypic gradients. Finally, we report LSGI as an open source R package https://github.com/qingnanl/LSGI.

## Results

### Overview of the LSGI framework

The main purpose of LSGI is to characterize spatial transcriptional gradient (STGs) of cells by answering three major questions: first, where does such gradient exist on the spatial map; second, what is the direction of the gradient; and third, what is the functional interpretation of the gradient (Figure 1A). To achieve this goal, LSGI by default employs NMF to factorize the collective gene expression profiles of all the cells or spots in a ST data into multiple programs (Figure 1B), including those delineating cellular compositions and those regulating cellular phenotypes. Through this step, cell loadings and gene loadings are calculated indicating the cell/spot-level activity of the programs and gene-level attribution to the programs, respectively. Since there is no prior information of the locality, linearity, and spatial mode (e.g., simple monotonical gradient or radial-like gradient) of the cells with STGs, we examine the spatial map with a slide-window strategy (Figure 1C), under which cells are grouped by spatial localizations in overlapping windows (Methods). We then fit linear models using spatial coordinates as predictors and cell NMF loadings as target, for every NMF program and every group of cells (Figure 1D). R-squared is used to evaluate goodness of fit with larger values indicating existence of STGs. The direction of a gradient is determined by the corresponding regression coefficients. These steps create a map containing the localization and direction of STGs as well as their assignment to one or more NMF programs. We then functionally annotate the programs by statistical methods (e.g., hypergeometric test) utilizing curated functional gene sets (Figure 1E, left). And investigate the spatial relationships of gradients assigned to different programs, or the spatial relationship of gradients to tumor-TME boundary in tumor ST datasets (Figure 1E, middle and right).

**Figure 1.**
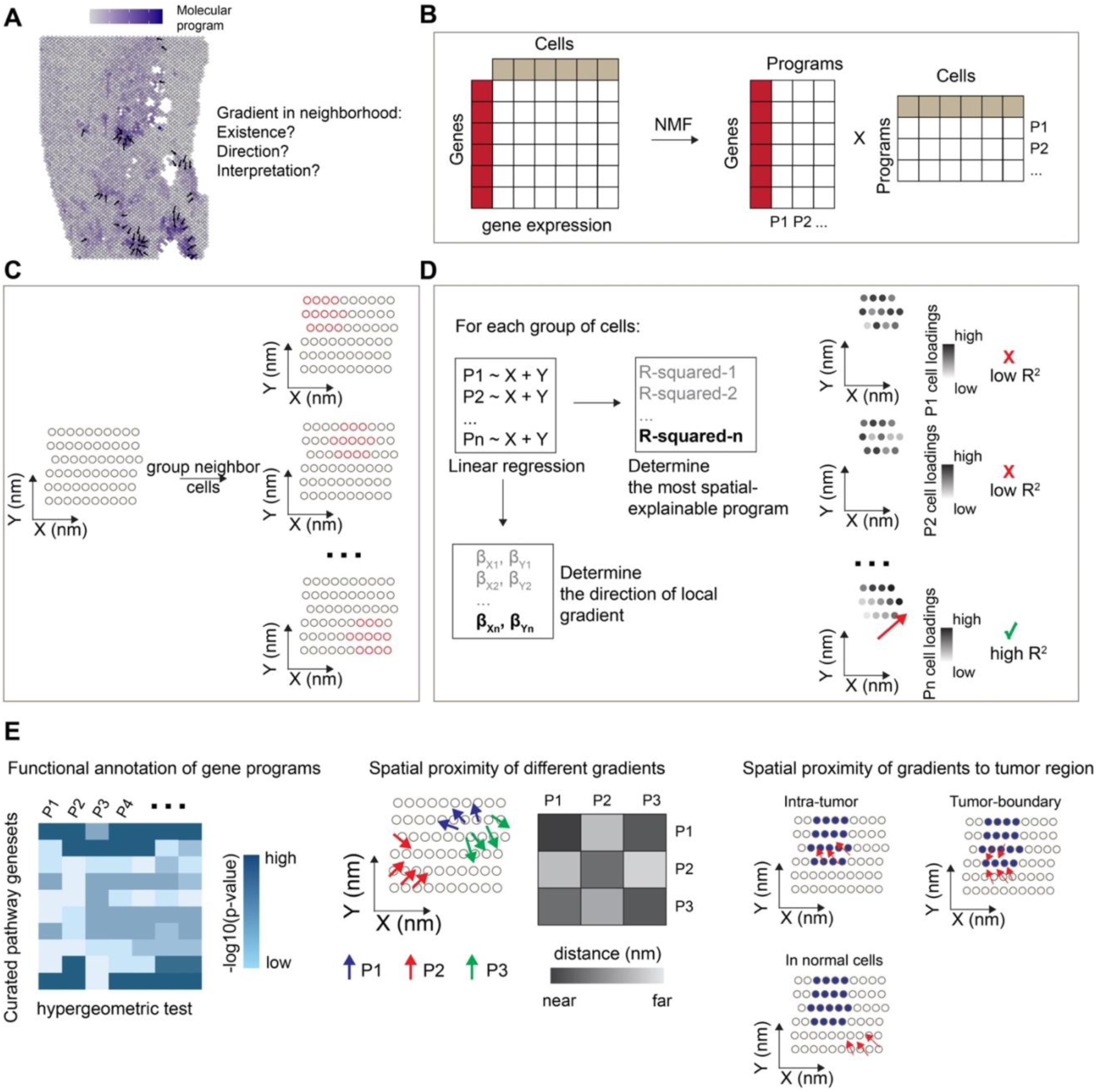
The LSGI framework and downstream analysis. A. Demonstrative plot showing existence of cell phenotypic gradient, summarized by some molecular programs, on a spatial map of tissue. Dark blue color demonstrates higher activity levels of the molecular program. Arrows indicate the direction of gradients. B. LSGI employs NMF to summarize the gene expression of cells into programs. C. LSGI partitions cells into small groups based on their spatial localizations. One cell can be assigned to multiple groups. D. Linear regression is performed in each spatial group of cells by fitting the loading of each NMF program with X and Y coordinate. R-squared is used to evaluate the performance of the regression, while the regression coefficient determines the direction of the local gradient. E. Downstream analysis on LSGI outputs: functional interpretation of gene programs (left), spatial proximity of different gradients (middle), and spatial proximity of gradients with other biological factors, such as the boundary of tumor core.

### LSGI reveals intratumoral, opposite-directed gradients of cell phenotypes in a GBM dataset

To investigate the power of LSGI in dissecting tissue heterogeneity, we first applied LSGI to a glioblastoma (GBM) ST dataset^22^ (UKF243_T_ST). In this experiment, we empirically identified STGs as those with R-squared higher than 0.6, and visualized them as arrows on the spatial map, colored by their assignment to different NMF programs (Figure 2A). We also highlighted tumor-harboring spots (through aneuploidy analysis) with grey circles to elucidate the spatial relationship between the STGs and tumor-TME boundaries (Methods, “cross-sample analysis in 87 tumor ST datasets: preprocessing and tumor region annotation”). We found that different NMF programs have distinct loading and STG distributions over the map and the patterns often coincide with the tumor-TME boundaries (arrows) (Supplementary Figure 1A-D).

**Figure 2.**
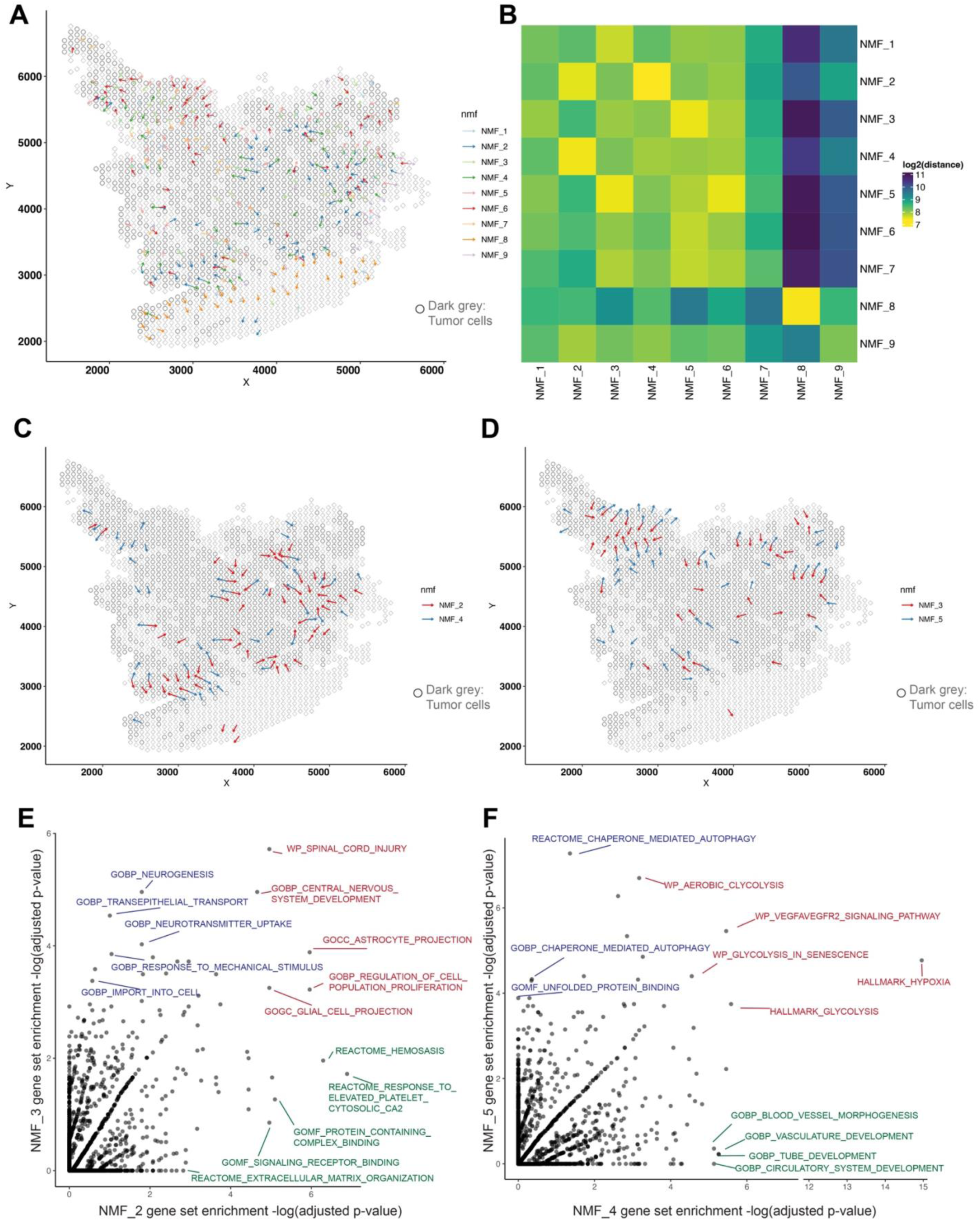
Application of LSGI on single ST dataset. A. Visualization of LSGI output on the spatial map (dataset: UKF243_T_ST from the RaviGBM study). Each rhombus represent a data spot (10x Visium technology) while the overlaying dark grey circles represent data spots characterized as tumor region. Each arrow indicate the presence of a gradient and the colors represent different NMF program of this gradient. Arrows directions indicate the direction of gradients. B. Spatial proximity of different gradients. The colors represent the log-transformed distance from the NMF program in a row to the program in a column. Here the distance is the real physical distance. Notice that this matrix is not symmetric (Methods). C-D. Visualization of the proximal NMF program pairs (C: NMF_2/4; D: NMF_3/5). Each arrow indicate the presence of a gradient and the colors represent different NMF program of this gradient. Arrows directions indicate the direction of gradients. The overlaying dark grey circles represent data spots characterized as tumor region. E. Comparison of pathway enrichment in top loading genes of NMF_2 and NMF_3. Each data point is a pathway and the two axes are the −log(adjusted p-value) for the hypergeometric test for enrichment. F. Comparison of pathway enrichment in top loading genes of NMF_4 and NMF_5. Each data point is a pathway and the two axes are the −log(adjusted p-value) for the hypergeometric test for enrichment.

We then quantified the mean physical distance between each type of the gradients, through which we noticed that some programs tend to colocalize, such as NMF_2 and NMF_4, or NMF_3 and NMF_5, etc. (Figure 2B, Methods). Interestingly, we observed that at multiple locations, the NMF_2 and the NMF_4 gradients colocalize yet pointing towards opposite directions, as if they repel against each other. Similar patterns were seen among the NMF_3 and NMF_5 gradients (Figure 2D).

To interpret these programs, we performed gene set enrichment analysis for each NMF program (based on top 50 genes by loading levels) through hypergeometric tests (Figure 2E). Interestingly, we found an enrichment of astrocyte and cell proliferation related terms in NMF_2 and NMF_3 (Supplementary Figure 2A and 2C) and the top genes include *SLC1A3* and *GFAP*, markers of the previously defined ‘AC-like’ tumor cell state ^23^. On the other hand, we found an enrichment of hypoxia related terms for NMF_4 and NMF_5 (Supplementary Figure 2B and 2D) with the top genes *VEGFA*, *NDRG1* and *ENO1*, markers of previously reported ‘MES-hypoxia’ tumor cell state^24^. Our findings are consistent with a previous study that cells with hypoxia and migration phenotypes display opposite orientations^22^. While the previous findings relied on knowing the genes a priori, our findings were *ab initio* from the ST data.

Besides the shared pathways, we also discovered differentially enriched pathways between each pair. For example, although NMF_2 and NMF_3 both had astrocyte-related terms, NMF_2 had several intercellular interaction terms such as extracellular matrix organization, signaling receptor binding, etc., while NMF_3 programs related to neuron functions such as neurogenesis and neurotransmitter uptake. Similarly, although NMF_4 and NMF_5 both had hypoxia and glycolysis terms, NMF_4 specifically had blood vessel development related terms, while NMF_5 autophagy related terms, highlighting different reactions upon hypoxia signals. Our findings also imply that different phenotypes marked by the paired NMF_2/3 and the NMF_4/5 programs are functionally coupled to each other.

### Systematic analysis of 87 tumor ST datasets with LSGI

To perform systematic tumor STG discovery, we further collected 87 ST datasets from 9 different studies (Table 1, Supplementary Table 1) including samples from a variety of tumor types. We performed LSGI independently on each sample (Figure 3A) and obtained at least one gradated NMF program greater than the empirical R-squared cutoff (>0.6) in 75 of the 87 datasets. From these NMF programs, we curated 19 meta-programs (Figure 3B) after merging similar programs using an approach published previously^25^ (Methods, “clustering NMF programs to meta-programs”). Some meta-programs consist of programs deriving from one tumor type or one study, while others were recurrent (Figure 3B, Supplementary Figure 3A-B). For each meta-program, we used the delta-Shannon entropy to quantify whether a large fraction of the meta-program was originated from a single tumor type or study (Figure 3C, Methods “calculation of compositional entropy”). Among the 19 meta-programs, 6 were identified as pan-cancer ones while the others were cancer type specific. We further annotated the meta-programs using functionally curated gene sets (Methods, Figure 3D) and visualized the loadings of the genes from assigned pathways in each program, grouped by the meta-program (Supplementary Figure 3C). Of particularly interest are the pan-cancer meta-programs related to EMT (epithelial mesenchymal transition), OXPHOS (oxidative phosphorylation), smooth muscle, extracellular matrix, and immune (MHC complex and B cell activation). The functional annotation of cancer-type specific meta-programs also showed consistency with prior knowledge, for example, MP-1 and MP-10 were related to keratinization, and they were solely originated from squamous cell carcinoma datasets. Moreover, MP-4 was related to hypoxia and was mostly originated from GBM datasets. Many of the terms have been previously reported in cancer single-cell studies, such as EMT, MHC, hypoxia, neurogenesis, etc. About 1/4 (90 out of 356) the programs were not clustered in meta-programs, highlighting the degree of intra-tumoral heterogeneity. Full information of the factors, their meta-program assignment, and the functional annotation of the meta-programs are reported in Supplementary Table 2-3.

**Figure 3.**
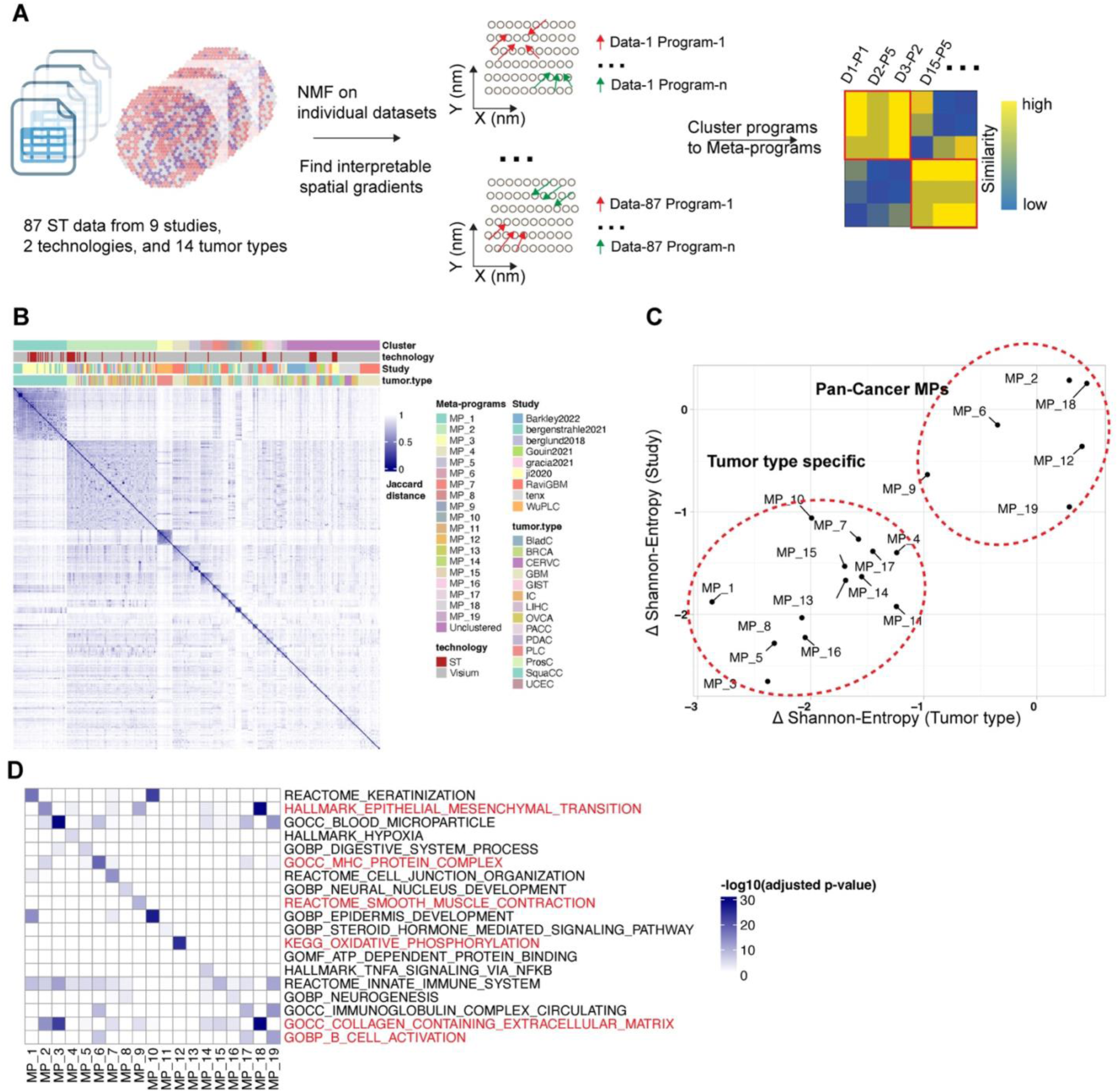
Cross-sample analysis of tumor datasets with LSGI. A. Schematic of the study design. LSGI was applied to each dataset separately and the NMFs were then integrated through clustering. B. Information of the 19 meta-programs. The heatmap showed the Jaccard distance between programs (using top 50 genes). Each program was labeled with the meta-program, technology, study and cancer type information. C. Study and tumor type specificity of each meta-program. The theoretical maximum Shannon entropy was calculated for each meta-program based on the tumor type and study label through averaging of random shuffling labels. These entropy quantifications were further subtracted by the real compositional Shannon entropy of the meta-program. D. Functional annotation of each meta-program. The pan-cancer meta-programs were highlighted with red labels of the annotation term.

**Table 1.** A brief summary of the 87 datasets used for cross-sample analysis with LSGI.

| Study Name | Number of datasets | Cancer type(s) |
| --- | --- | --- |
| Barkley2022 | 10 | BRCA (breast cancer), GIST (gastrointestinal stromal tumor), LIHC (liver hepatocellular carcinoma), OVCA (Ovarian cancer), PDAC (pancreatic ductal adenocarcinoma), UCEC (uterine corpus endometrial carcinoma) |
| Bergenstrahle2021 | 8 | IC (intestine cancer), SquaCC (squamous cell carcinoma) |
| Berglund2018 | 14 | ProsC (prostate cancer) |
| WuPLC | 7 | PLC (primary liver carcinoma) |
| Gouin2021 | 4 | BladC (bladder cancer) |
| Gracia2021 | 4 | OVCA |
| Ji2020 | 16 | SquaCC |
| tenx | 6 | BRCA, CERVC (cervical cancer), IC, OVCA, PACC (prostate cancer, adenocarcinoma with invasive carcinoma), ProsC |
| RaviGBM | 18 | GBM (glioblastoma) |

We then sought to investigate whether the spatial locations of the STGs can inform tumor-TME tissue architecture. We performed the analysis in the following steps: First, we annotated tumor spots with aneuploid copy number profiles using CopyKat (Methods, Supplementary Figure 4A); Second, with the tumor/normal spot annotation, we calculated a tumor ratio in each sliding window (Methods, “Calculation of tumor ratios for each local gradient”, Supplementary Figure 4B); Third, for a given STG (associated with one NMF program), we collect the tumor ratio in its constituent sliding windows and calculate the average. Intuitively, a low average tumor ratio indicates that the STG tends to appear within normal tissue regions, while a high ratio indicates that the STG tends to appear within tumoral regions. We demonstrate three examples representing low, medium, and high average tumor ratios, respectively (Figure 4A-C). In each panel, the overlaying dark grey circles represent data spots characterized as tumor region by CopyKat (Methods). Indeed, we found lower average tumor ratios indicated association to normal regions while higher values indicated association to tumor regions, and medium values to the tumor-TME boundary. We clustered the average tumor ratios for all the (356) programs and categorized them into three tumor ratio clusters (TRCs, Supplementary Figure 5A-B), and we noticed differential proportion of TRCs among meta-programs (Supplementary Figure 5C). Here, we demonstrate a few examples of MP_14, annotated as TNFA signaling via NFKB. All four programs in this meta-program were from GBM datasets and three of them (UKF243_T_ST, NMF_6; UKF260_T_ST, NMF6; UKF255_T_ST, NMF_3) were enriched in intratumoral region (all belonged to TRC Cluster 3). We confirmed their localizations in intratumoral regions through visualization of the gradients and representative genes such as *FOS, CD44, DUSP1, ZFP36*, etc. (Figure 4D-E, Supplementary Figure 6A-D). The activation of TNF-NFKB axis has already been revealed in several tumor types including the GBM^21,26^, while here, through a systematic analysis, we unraveled its association to spatial intratumoral heterogeneity, with consistency in several patient samples. Finally, all the LSGI outputs for these tumor datasets were made accessible (https://zenodo.org/records/10626940) and sample codes and detailed tutorials were available to the community to freely explore and visualize the data.

**Figure 4.**
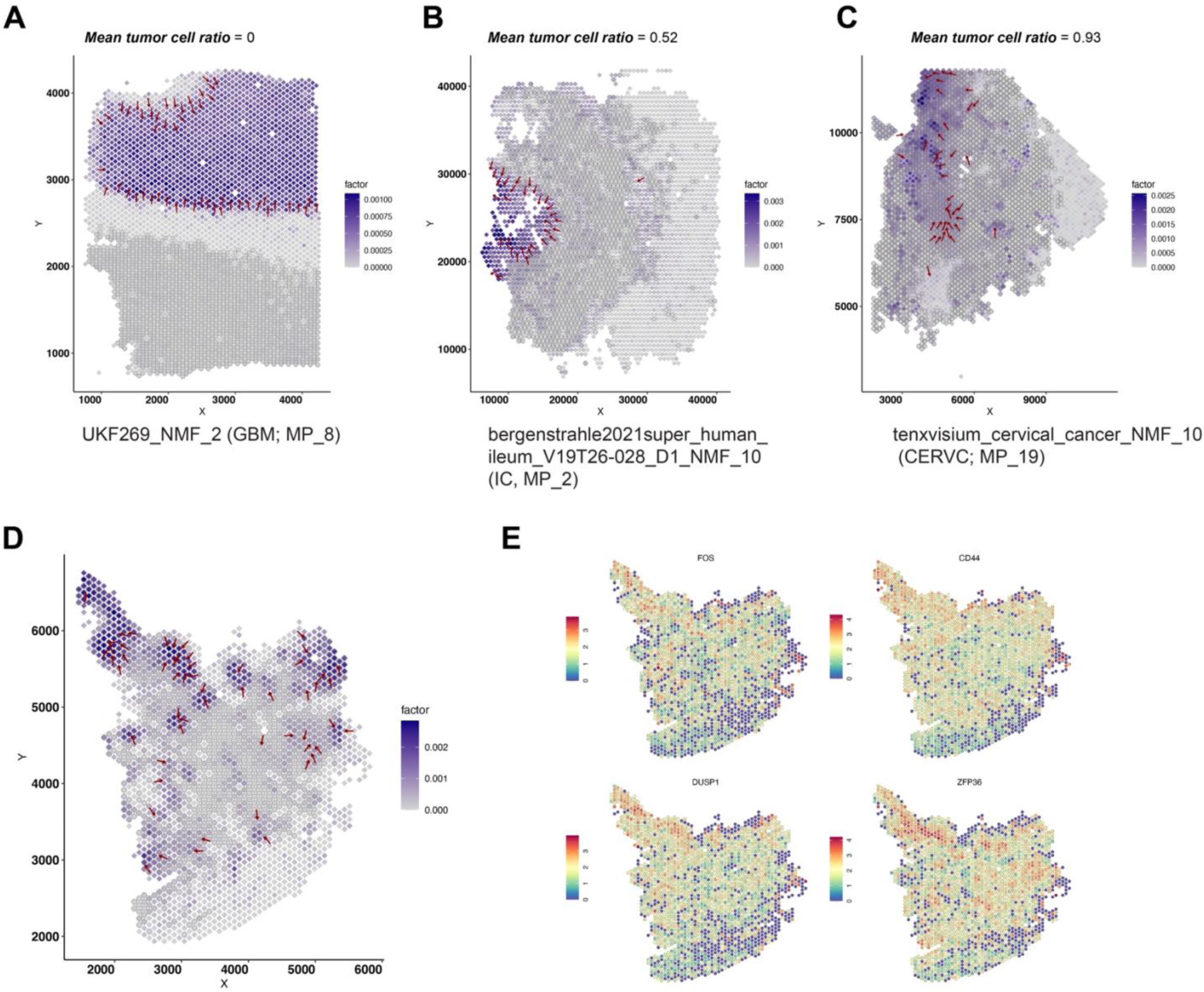
Spatial relationship between gradients and tumor boundary. A-C. Examples of NMF programs with different mean tumor cell ratios. The information of the program, its tumor type, and its meta-program assignment was labeled under each panel. Red arrows marked the presence and direction of the gradient. For each panel, each rhombus represent a data spot while the overlaying dark grey circles represent data spots characterized as tumor region. Color of the rhombus represent the loading of the NMF program. D. The gradient direction and original cell loadings of NMF_6 (UKF243_T_ST) on the spatial map. The overlaying dark grey circles represent data spots characterized as tumor region. E. The spatial expression of representative genes in NMF_6 (UKF243_T_ST). Warmer colors (red) indicate higher expression levels.

## Discussion

In this study, we introduced a simple, flexible yet highly interpretable strategy, LSGI, for discovering spatial transcriptomic gradients in a ST data. Given the uncertainty of the existence and spatial variation of STGs, we employed a divide-and-conquer strategy by calculating local linear gradients in sliding windows, which collectively produce a STG map across the tissue. We demonstrated the utility of LSGI for both in-depth, single dataset analysis and cross-sample meta-analysis using 87 tumor ST datasets. Without any prior knowledge, from merely 87 samples, LSGI was able to identify gene expression programs consistent with prior cancer studies and discover patterns indicative of spatial transcriptional heterogeneity on each tissue slide, providing novel functional annotations and insights that would otherwise be missed by the current ST data analysis practices^27^ (cell clustering and annotation, spatial niche assignment, spatially variable gene analysis, etc.), or manual, image-based annotations. Compared to the approaches that summarize spatial data into static spatial domains, we showed that spatial gradient approaches were capable of deconvoluting the tumor state dynamics in the spatial context. As we showed, some tumor regions could be associated with different types of phenotypical gradient at various levels, while assignment of such regions to a single niche/domain would likely lose such dynamical view. Moreover, the development of LSGI also enables association analysis between STGs and pathological/morphological annotations to deepen our knowledge of molecular pathology.

We demonstrated the utility of LSGI on sample datasets generated by 10X Visium and an early version of spatial transcriptomics^28^, as they have whole-transcriptomic coverage thus enabling unbiased functional interpretation of NMF programs. The spatial resolution of those technologies, however, can limit the power of discovery and confound the result due to cell admixing in spots. This is a limitation of the data, not of LSGI. The LSGI framework can be applied agnostically to technologies, as the only required inputs are spatial coordinates and gene expression levels. As single-cell resolution whole transcriptomic ST technologies^29^ becomes increasingly available, we expect a relatively straightforward adaption of LSGI into new technologies. Lastly, although not demonstrated in this study, LSGI can easily fit three-dimensional ST data analysis through adding an additional ‘Z’ coordinate to the linear regression step.

Noticeably, several other methods^30–32^ also aimed to detect gradated signals in ST data. While these methods focus largely on inference of global spatiotemporal trends from continuous gene expression data, LSGI focuses on detecting interpretable, phenotypically salient gradients factorizable by NMF. We propose that caution needs to be taken when attempting to use all cells/spots to infer a ‘global’ gradient, because when no biologically meaningful gradients are present (for instance, distinct cell types mixed together in some regions of complex tissues), trajectory inference method may overfit the data. To this end, LSGI benefits from its design that the existence of each local gradient is assessed by how well the local linear model fits the data. In practice, we found that many regions on ST data do not have salient gradients of any programs. In the meta-analysis approach of this study, 12 of the 87 tumor samples had no salient gradient identified. Finally, LSGI is benefited from its utilization of NMF to extract transcriptional phenotypes from the expression matrix because NMF has been shown capable of capturing biological signals^33^ and have been widely applied in single-cell or ST studies^25,34–38^. The employment of NMF not only enhances the interpretability of LSGI, but also allows effective cross-sample comparison of the programs, as was previously reported, which laid the basis of our meta-analysis approach to find recurrent STGs in different tumors. Thus, we believe that LSGI serves as a powerful and complementary approach to the other methods targeting alternative scopes and resolution.

Although by default, LSGI uses basic NMF to factorize gene expression programs, the LSGI framework is flexible and can accommodate variants of NMF methods, such as cNMF^39^, iNMF^40^, and jNMF^41^, or other types of cell phenotypic quantification, such as pathway activity measurements^42,43^ to calculate their spatial gradients. We do recommend using NMF if no assumptions were made, and if the users required a systematic, unbiased analysis. For datasets generated with targeted ST technologies^44,45^, we suggest that users be cautious in annotating the NMF programs as the gene set (panel) may be biased towards some pathways due to biased selection of genes.

## Methods

### The LSGI framework

The main LSGI framework starts from clustering spots (or cells for datasets with single-cell resolution; we would refer to this unit as cell in the following description for simplicity) into small groups solely based on their localization. The number of groups *P* is controlled by a parameter *S*, that 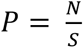, where *N* is the total number of cells. By default, *S* is set to 5, while the group size *Q* is set to 25. Thus, each cell is included in 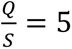 groups on average. The selection of *Q* controls the resolution of the gradient detection. Setting a smaller *Q* would let the LSGI program examine linear gradients within smaller window sizes (higher resolution) while also has the risk of reduced robustness to noise due to smaller sample sizes in multivariate regression. We also require such groups of cells to be tiling for reducing unwanted effects of arbitrarily determining the groups by suggesting a smaller *S* than *Q*. To achieve such tiling, we used the ‘balanced_clustering function in the ‘anticlust’ R package^46^ to cluster cells into *P* groups based on the spatial coordinates and determine a grid point at the center of each cluster. We then search for the *Q* nearest neighbors among cells to each grid point, based on Euclidian distance, thus forming the groups.

By default, LSGI take NMF embeddings of cells as the input. The NMF step is not incorporated in the LSGI framework as many NMF implementation have been reported and we want to offer this flexibility to users. All the NMF step involved in this work used the NMF implementation of the singlet R package^47^.

With the group and NMF information, a linear regression is performed for each NMF program in each group: 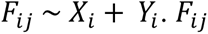 is the loading of the cells from the *i*th group of the *j*th NMF program. *X*_*i*_ and *Y*_*i*_ are the spatial coordinates of the *i*th group of the cells. The regression coefficients *β_Xij_* and *β_Yij_* determine the most likely gradient direction of this program *j* in the group *i*, while the *R*^2^ of this regression represents the largest explanatory capability of spatial effects on the cell loadings of program *j*. Such processes are performed iteratively for all NMF programs and all cell groups. Although *R*^2^ has a clear statistical meaning, the selection of its threshold could be empirical given different contexts. In this study, we only treated the cases where *R*^2^ ≥ 0.6 as valid gradients and these were retained for further analysis. As *R*^2^ equals to 0.5 often treated as a moderate goodness of fit and our rationale was to call the gradient where a slightly higher proportion of the molecular signal (NMF loadings) explained by the spatial localization. Additionally, we only retain programs with gradients in at least 5% of total grid points. For the ‘arrow’ visualization (such as Figure 2A), the arrow directions are pointing to increased program signals (such as Figure 4D). Please note that it is possible that one group of cells can have different gradients assigned to different NMF programs (usually gradated to different directions).

Furthermore, the LSGI package offers a strategy to estimate the overall distance between two types of gradients (as is shown in Figure 2B): Overall distance from a gradient *A* to gradient *B*, *D*(*F*_*A*_, *F*_*B*_), is calculated: 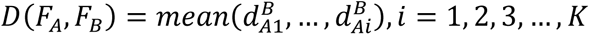; 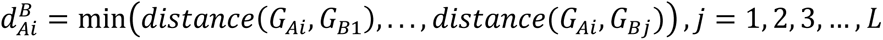 K is the number of grid points (*G*) with gradated program *A*, L is the number of grid points with gradated program *B*, *distance* here is Euclidean distance. In short, for each grid point *i* with program *A* (*G_Ai_*), we find the closest grid point with program *B* and record that distance (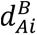). We then use the mean of this distance of all grids with program *A* as an overall evaluation of closeness from gradient *A* to gradient *B*. Please note here *D*(*F*_*A*_, *F_B_*) ≠ *D(F_B_*, *F_A_*).

LSGI is an efficient program that the main gradient inference step takes less than 1minute for each dataset in our practice (roughly 3000-8000 spots per dataset, 16 GB RAM MacOS laptop). The LSGI R package has been tested on MacOS (Ventura 13.6), Windows (Windows 11), and Linux (Redhat Enterprise) systems.

### Cross-sample analysis in 87 tumor ST datasets: Preprocessing and tumor region annotation

All the ST datasets were curated and converted to Seurat objects and were preprocessed following the SeuratV4^48^ workflow, including normalization, scaling, dimensionality reduction (with PCA) and clustering, with default parameters. NMF was performed with the singlet R package^47^, scanning the number of factors *k* with a range from 6 to 10. The final *k* value was decided by cross-validation implemented in the same package. Tumor regions of ST datasets were inferred using CopyKat^49^ with automatically determined normal cell references. Given the prevalence of immune cells in tumor samples, we sought to use immune cells as the normal cell references (Supplementary Figure 4A). We quantified the expression of a set of immune related genes (CD3E, CD8A, GZMK, CD4, CCR7, GZMB, FCER1G, LHDB, DUSP2, IL7R, S100A4) at the single-cell level, and then treat the cluster (from Seurat) with the most top immune-related cells (top 100 cells with highest immune gene expression) as the normal cell reference. The other parameters were default for CopyKat.

### Calculation of tumor ratios for each local gradient

With the annotation of tumor spots, we could obtain the tumor ratio for each grid point (Supplementary Figure 4B). For a gradated NMF program of a dataset, we collected the tumor ratio for all grid points where it showed gradient (*R*^2^ ≥ 0.6) and used the average tumor ratio to concisely summarize the spatial relationship between that program and tumor core, normal tissue, or tumor-TME boundary. To cluster programs into different tumor ratio clusters (TRCs), equal-weighted one-dimensional K-nearest neighbors clustering were applied (Supplementary Figure 5A-B).

### Clustering NMF programs to meta-programs

We then applied LSGI separately on each dataset and combined the output for an integrative analysis. We retained only the gradated NMF programs in at least 5% of the total grid points for each dataset. We then clustered the remaining NMF programs following a previously reported approach^25^. Briefly, each cluster of the programs started from a founder program that having the most high-overlapping cases (over 20 overlapping genes among top 50 with highest loadings) with other programs (at least two other programs). The founder program would then be clustered with the program with highest overlapping genes (and at least 20 overlapping genes), and this meta-program will be assigned a 50-gene signature based on their appearance in the top 50 of each program and their loadings of the original NMF program. The cluster would further grow iteratively following such rules until no programs can be merged into it. Such processes would then start again in the rest of the programs until no founder programs could be identified, and the program left would be assigned to the ‘Unclustered’ group. Thus, each meta-program (except the ‘Unclustered’) was be summarized as a 50-gene signature which facilitated the functional annotation of the meta-program.

### Calculation of compositional entropy

To quantify whether a meta-program was formed by programs from specific study or tumor types, we calculated the delta-Shannon entropy for each meta-program. The Shannon entropy for each program is: 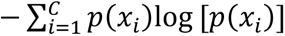. *C* is the number of categories (tumor type or study), while *p(x_i_*) here is the fraction of meta-program originated from the *i* th category. We then shuffled the category labels randomly and calculated the simulated random entropy and subtracted the average random entropy (10 times simulation) from the real entropy to obtain delta-Shannon entropy. Such measurement reflect how likely a meta-program is composed of programs from different categories with the same probability.

### Functional annotation of NMF programs and MPs

For functional annotation of NMF programs, we tested the enrichment of functional gene sets in the top 50 genes in each program with highest loadings, while for meta-programs, the 50-gene signatures were directly used. The hypergeometric tests were performed with the R package hypeR^50^. Several functional gene sets were combined as the input: Gene Ontology^51^ (Biological Process, Molecular Function, and Cellular Component), MSigDB Hallmarks^52^, and Canonical Pathways from MSigDB C2 collection^53^. To decide the annotation of meta-programs, we first reduce the hypergeometric test results to top 40 gene set for each meta-program based on adjusted p-value (false discover rate adjusted), and further reduce the result to top 5 based on cross-program specificity. The specificity score for gene set *i* of meta-program *p* is calculated by 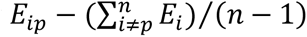. Here *E_ip_* is the negative log-transformed adjusted p-value for gene set *i* enrichment of meta-program *p* (hypergeometric test). Full functional annotation results are available in Supplementary Table 3.

## Data Availability

A summary of ST datasets is included in Supplementary Table 1. Most of the datasets were downloaded from the SODB curation (Barkley2022^34^, Bergenstrahle2021^54^, Berglund2018^55^, Gouin2021^56^, Gracia2021^57^, Ji2020^58^). 10x datasets were downloaded from the 10x Genomics website. WuPLC^18^ datasets were downloaded from https://ngdc.cncb.ac.cn/gsa-human/browse/HRA000437. RaviGBM^22^ datasets were downloaded from https://datadryad.org/stash/dataset/doi:10.5061/dryad.h70rxwdmj.

The LSGI processed data (87 tumor datasets) are available in https://zenodo.org/records/10626940. Sample analysis code (https://zenodo.org/records/10626940/files/LSGI-annotation-and-visualization-demo.html?download=1) are available for users to visualize and explore the data.

## Code Availability

LSGI is an open source R package hosted in GitHub: https://github.com/qingnanl/LSGI. The code used for analyzing the tumor ST data (preprocessing, running LSGI, and downstream analysis) is available at https://github.com/qingnanl/LSGI_manuscript_code/.

## Author Contributions

Q.L. and K.C. conceived the study. Q.L implemented the software, analyzed data, and prepared figures. L.S. and C.H. provided pathology insights of the method and contributed to the result interpretation. Q.L and K.C. drafted the manuscript with input from all. K.C. supervised the project.

## Competing Interests

The authors declare no competing interests.

## Acknowledgements

This project has been made possible in part by grant U01CA247760 to KC, and Human Cell Atlas Genetic Ancestry Network Grant CZF 2021-239847 to KC from the Chan Zuckerberg Initiative DAF, an advised fund of Silicon Valley Community Foundation. This project was also partially supported by the MD Anderson Moonshot programs. We also thank Nejla Lermi and Vakul Mohanty for valuable discussions.

---

**Supplementary Figure 1.**
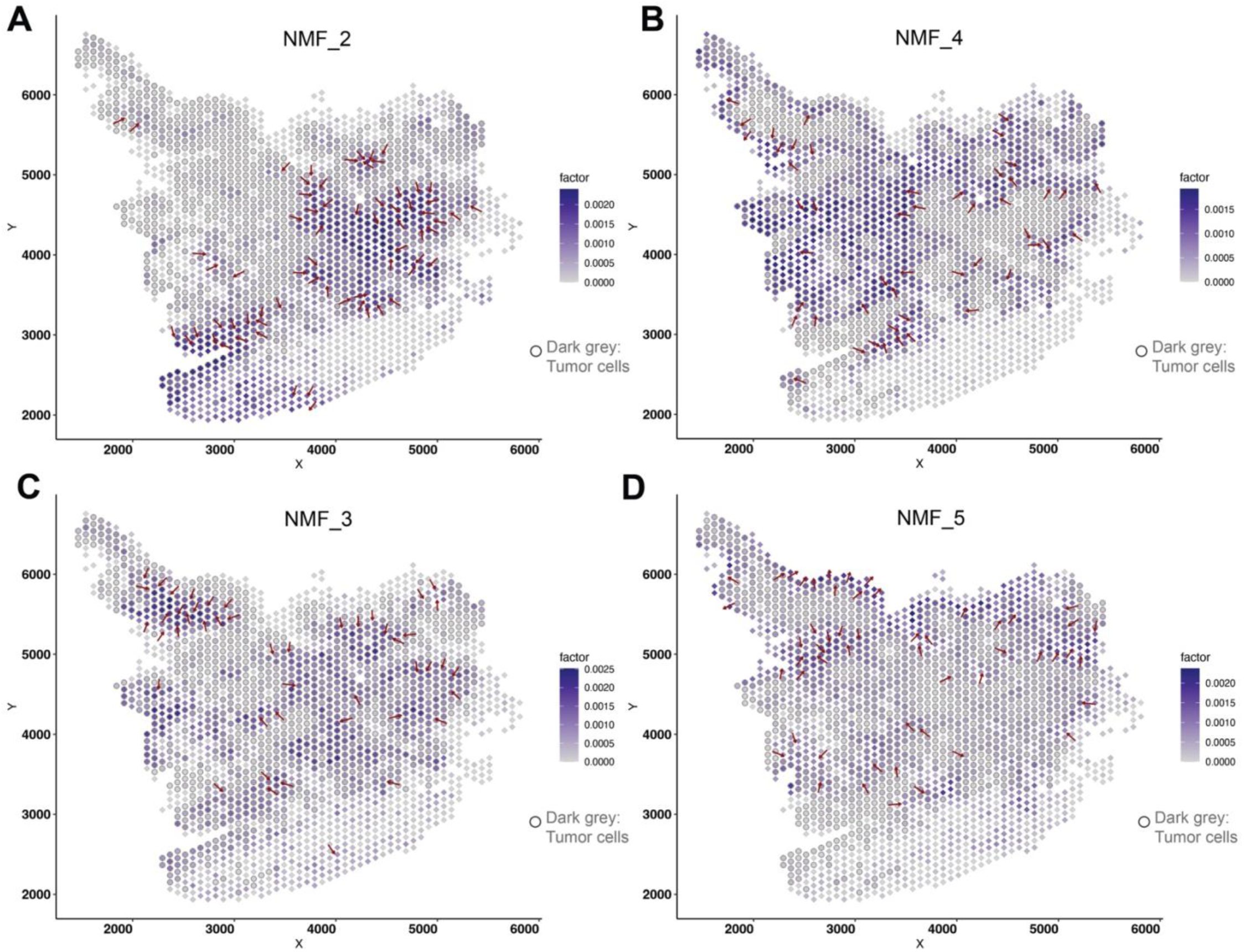
A-D. Demonstration of the gradient direction and original cell loadings of NMF_2 (A), NMF_4 (B), NMF_3 (C), and NMF_5 (D) on the spatial map. The overlaying dark grey circles represent data spots characterized as tumor region.

**Supplementary Figure 2.**
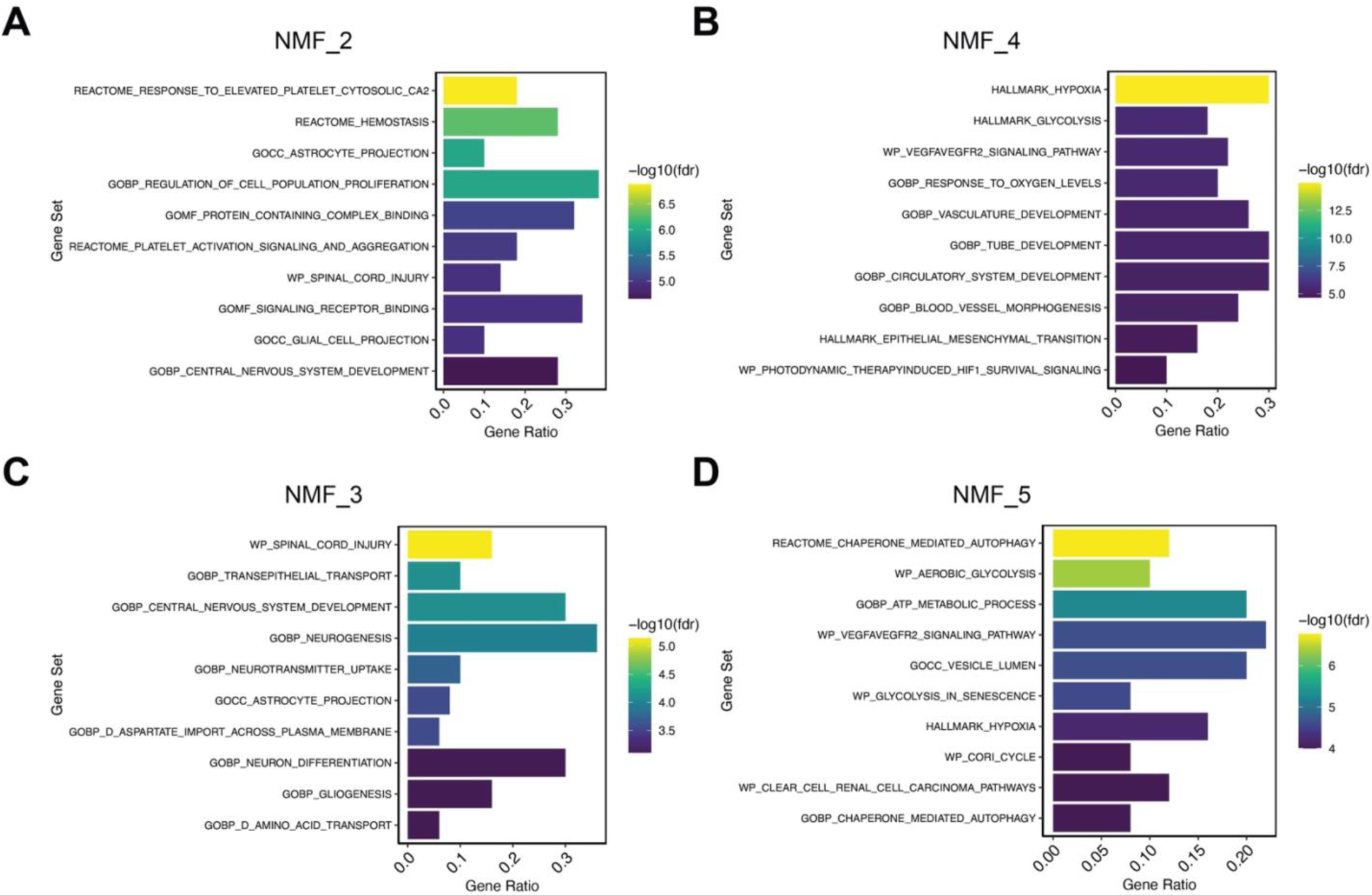
A-D. Functional enrichment of top genes in NMF_2 (A), NMF_4 (B), NMF_3 (C), and NMF_5 (D). Bar-plots showed the ratio of pathway genes found in input gene sets (top 50 genes in each NMF program) and were colored by the adjusted p-value (false discovery rate, FDR) of hypergeometric test.

**Supplementary Figure 3.**
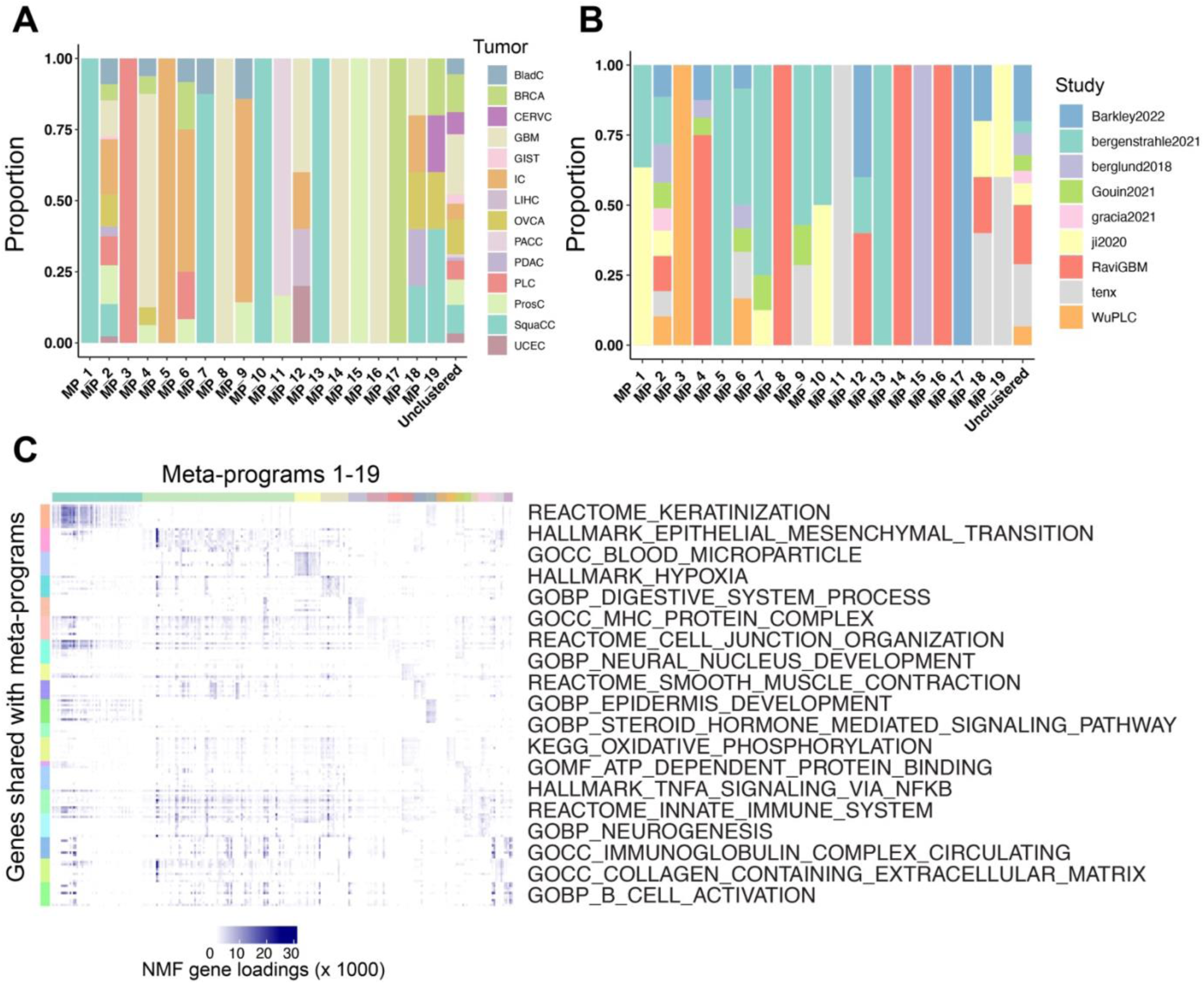
Information of meta-program composition and functional annotations. A-B. Proportion of originated tumor types and study of NMF programs in each meta-program. C. Loadings of the assigned functional gene set members in each NMF program grouped by meta-program.

**Supplementary Figure 4.**
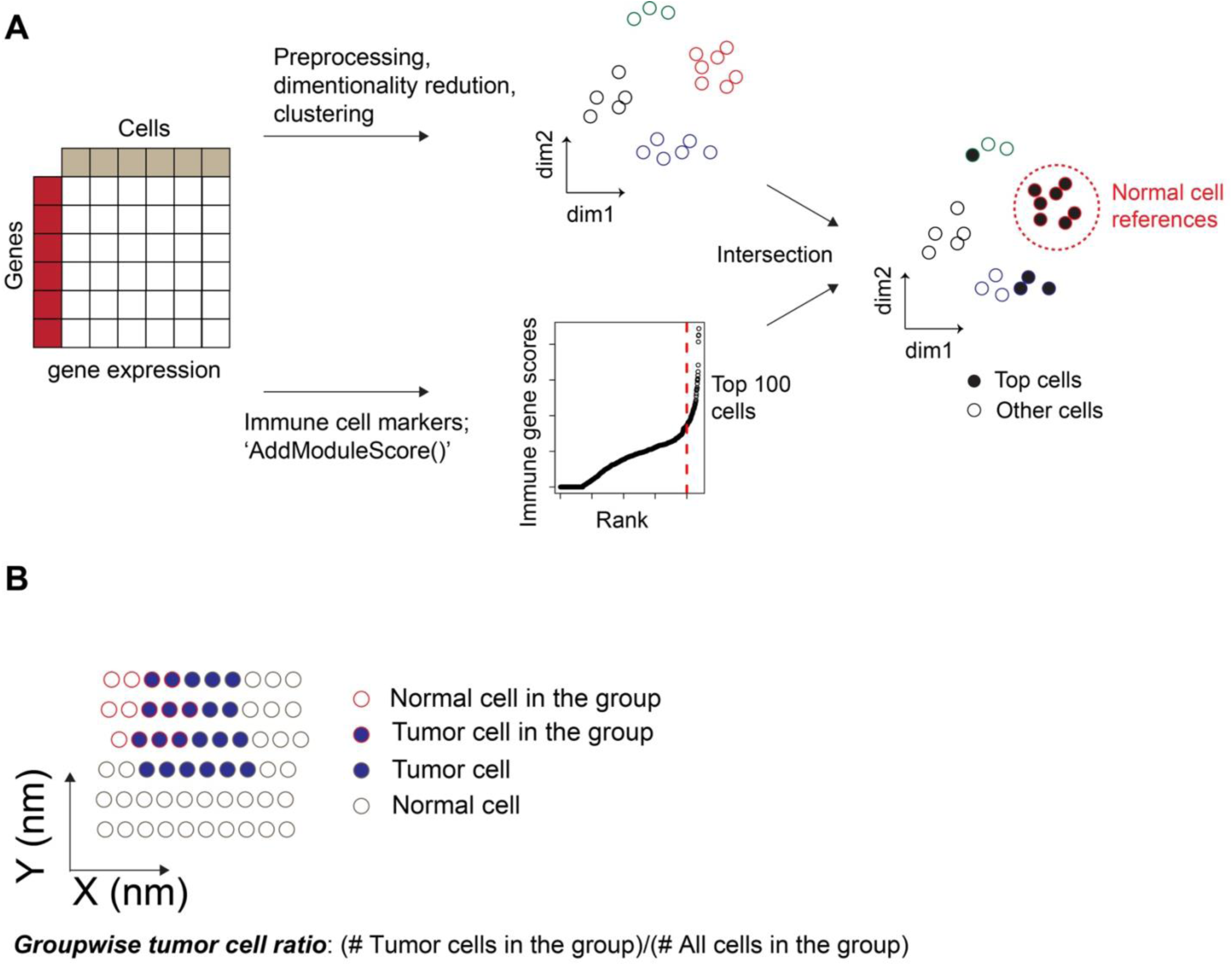
Strategy of automatically detection normal cell references and calculate tumor cell ratio in each local group. A. Strategy to infer immune cell clusters for annotating tumor regions with CopyKat. B. Calculation of groupwise tumor cell ratio. Red circles represent cells in a local group while grey circles are other cells. Dark blue labels tumor cells. For each group, the tumor ratio equals to the number of tumor cells in this group divided by the number of all cells in the group.

**Supplementary Figure 5.**
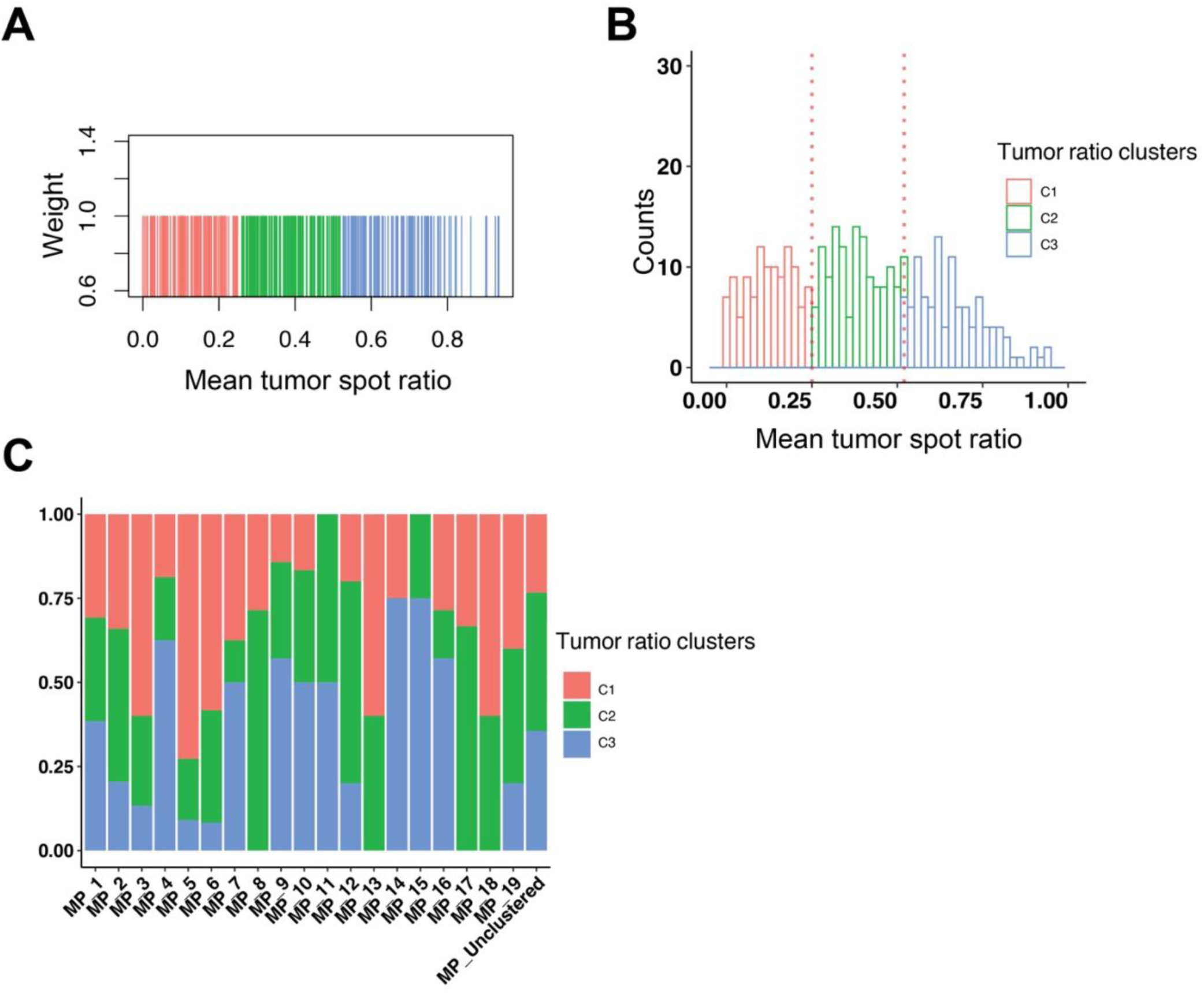
Cluster programs based on tumor spot ratios. A. Using one-dimensional equal-weighted KNN clustering of tumor cell ratios to form three TRCs. B. The distribution of tumor cell ratios among programs (right). Colors indicate the identity of three TRCs. C. Proportion of programs clustered to the three TRCs in each meta-program and the ‘unclustered’ programs.

**Supplementary Figure 6.**
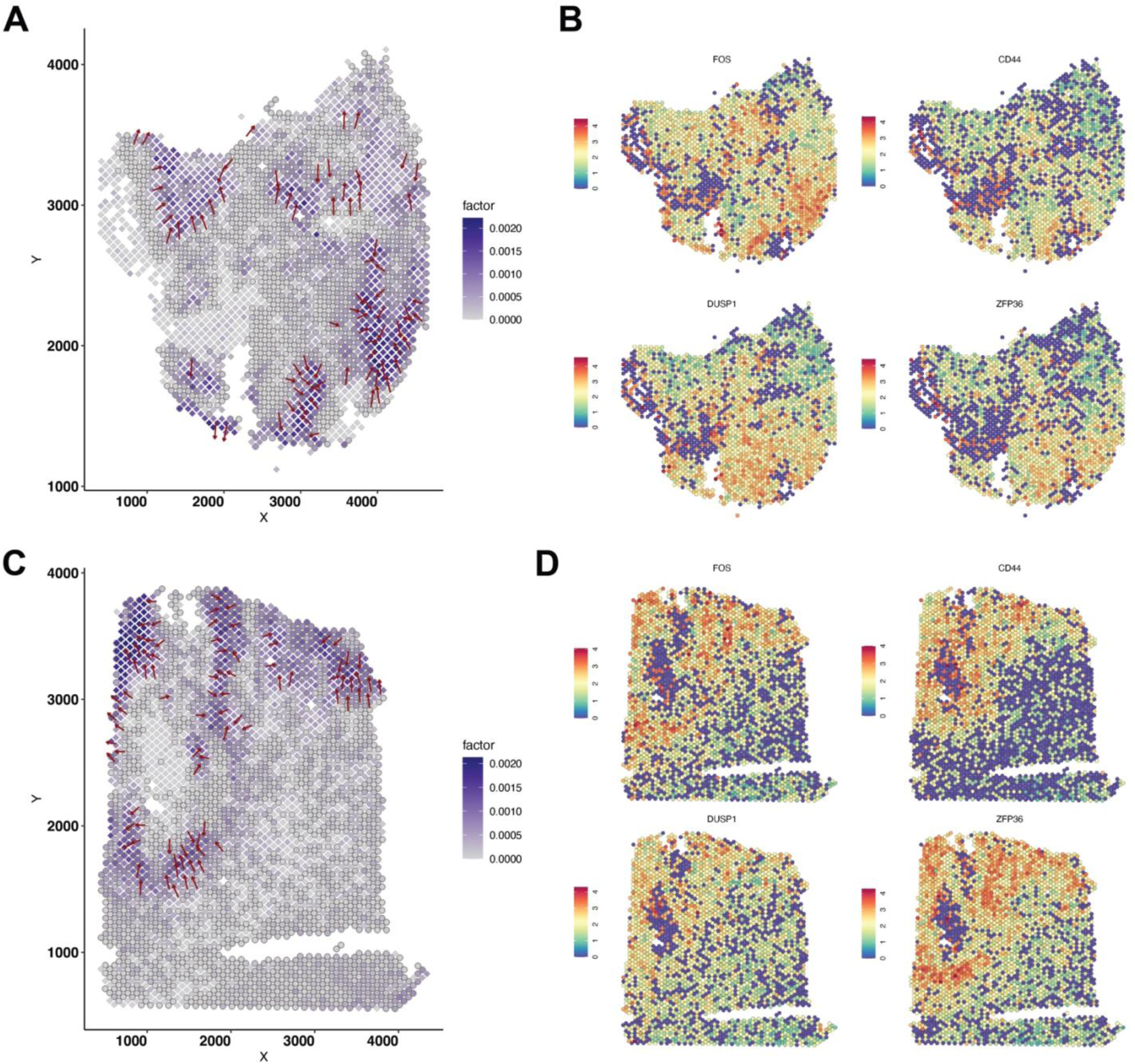
A. The gradient direction and original cell loadings of NMF_3 (UKF55_T_ST) on the spatial map. The overlaying dark grey circles represent data spots characterized as tumor region. B. The spatial expression of representative genes in NMF_3 (UKF255_T_ST). Warmer colors (red) indicate higher expression levels. C. The gradient direction and original cell loadings of NMF_6 (UKF260_T_ST) on the spatial map. The overlaying dark grey circles represent data spots characterized as tumor region. D. The spatial expression of representative genes in NMF_6 (UKF260_T_ST). Warmer colors (red) indicate higher expression levels.

## Notes

### Competing Interest Statement

The authors have declared no competing interest.

